# Unraveling cell differentiation mechanisms through topological exploration of single-cell developmental trajectories

**DOI:** 10.1101/2023.07.28.551057

**Authors:** Emanuel Flores-Bautista, Matt Thomson

## Abstract

Understanding the circuits that control cell differentiation is a fundamental problem in developmental biology. Single-cell RNA sequencing has emerged as a powerful tool for investigating this problem. However, the reconstruction of developmental trajectories is based on the assumption that cell states traverse a tree-like structure, which may bias our understanding of critical developmental mechanisms. To address this limitation, we developed a framework, TopGen, that enables identifying topological signatures of functional biological circuits as persistent homology groups in transcriptome space. First, we show that TopGen can identify genetic drivers of topological structures in simulated datasets. We then applied our approach to more than ten single-cell developmental atlases and found that topological transcriptome spaces are predominantly path-connected and only sometimes simply connected. Finally, we applied TopGen to analyze gene expression patterns in topological loops representing stem-like, transdifferentiation, and convergent cell circuits, found in *C. elegans, H. vulgaris*, and *N. vectensis*, respectively. Our results show that some essential differentiation mechanisms use non-trivial topological motifs, and that these motifs can be conserved in a cell-type–specific manner. Thus, our approach to studying the topological properties of developmental transcriptome atlases opens new possibilities for understanding cell development and differentiation.

## 1 Introduction

A major goal of single-cell genomics is to define properties of gene regulatory circuits from samples of molecular data such as the transcriptome and epigenome. Single-cell profiling has represented a powerful tool for investigating developmental processes in order to unravel regulatory control of cell differentiation at the transcriptional level [(1), (2), (3), (4), (5), (6), (7), (8), (9), (10)]. One of the central hypotheses in developmental biology is that because cell lineages consist of bifurcation events, cell states traverse a tree-like branching structure in gene expression space [(1), (11), (12)]; we will refer to this notion as the *tree hypothesis* (Fig 1 A). Mathematically, a branching tree can be characterized by its topology as a path-connected set of points in gene expression space that lacks holes or cycles and can, therefore, be contracted continuously to a point. In fact, formal tools from the field of algebraic topology can be applied to ask whether developmental cell trajectories in gene expression space, in fact, generate a contractible tree-like structure. The Betti numbers quantify the overall shape of a topological space. Formally, viewing the transcriptome as a topological space, the Betti numbers under the tree hypothesis become *β*_0_ = 1, indicating that developmental trajectories are path-connected and *β*_*i*_ = 0 for all *i >* 0, indicating that gene expression trajectories are cycle-free during development.

**Figure 1:**
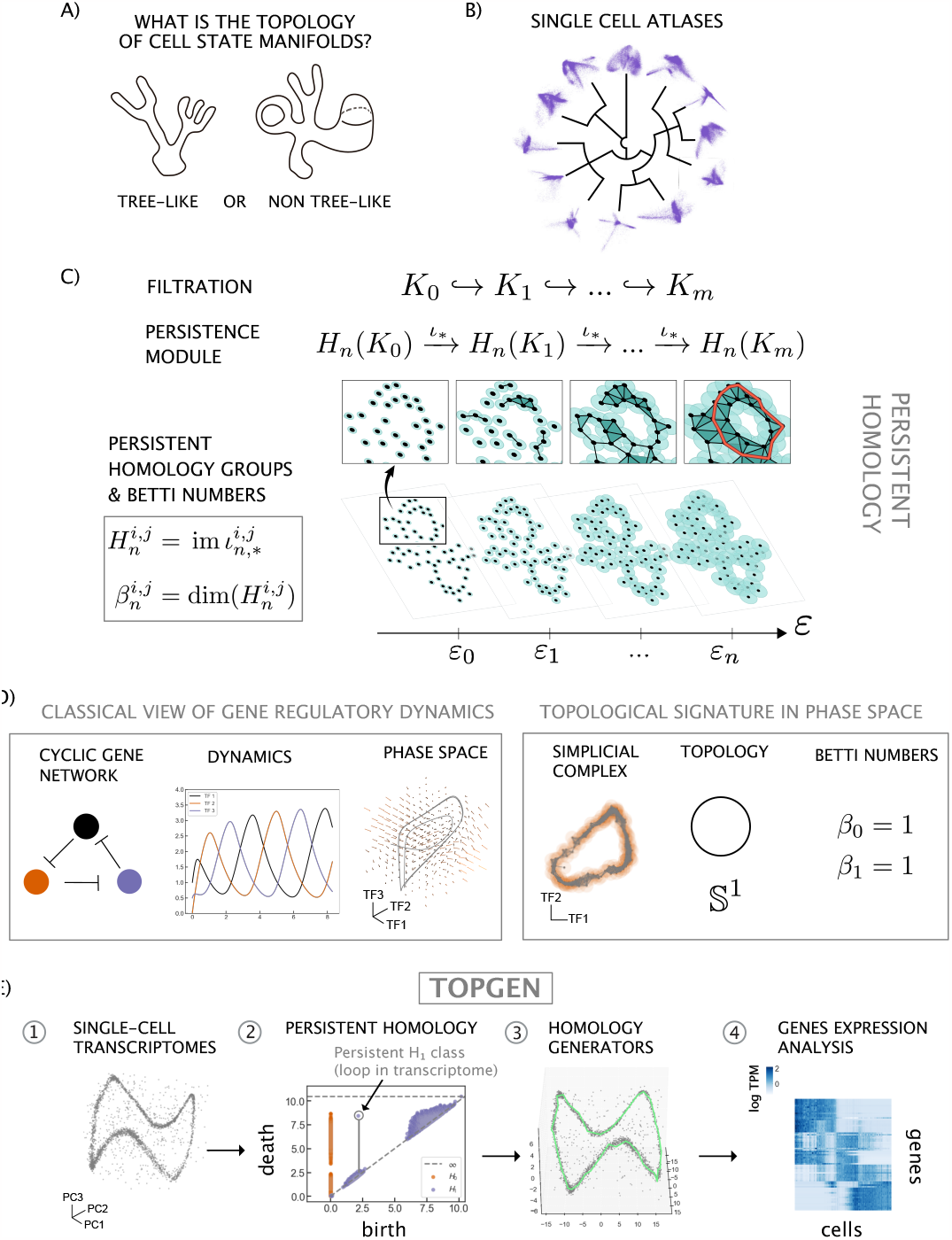
Discovering cellular differentiation dynamics using TopGen. (A) The current view of the topology of the transcriptome is that cells form a tree during fate commitment, which we refer to as the *tree hypothesis*. Cyclic or converging cell fates could break the tree hypothesis. (B) Phylogenetic tree diagram of survey organisms. We performed a topological survey of transcriptome atlases during early development using persistent homology. (C) Persistent homology (PH) is a robust method to identify topological signatures from noisy samples of a manifold. PH works by recording when topological features appear and decay (Methods). A 1-homology class is born in the rightmost panel. (D) Intuition of an *H*_1_ topological signature using a genetic oscillator. Dynamical systems view of gene regulatory control (left) has a corresponding topological interpretation (right) that can be formalized using homology groups and Betti numbers. (E) Workflow of TopGen to identify genetic drivers in single-cell transcriptome data.

Until recently, we lacked sufficient data to rigorously test the tree hypothesis during development (Fig 1 A). New single-cell atlases of organismal development provide trajectories across developmental time in different organisms. However, the complexity and high dimensionality of transcriptome spaces have rendered the rigorous testing of this hypothesis prohibitively hard. In congruence, analyses of developmental data manifolds either assume this hypothesis using *ad hoc* methods [(1), (13)] or use non-linear dimensionality reduction to get a global view of the space [(2), (4)]. However, it has been recently discovered that widely adopted methods like tSNE and UMAP can potentially distort the shape of data, for example by increasing the number of clusters (14). The use of these tools could thus bias and obscure the interpretation of the geometric and topological properties of cell trajectories in gene expression space and hide critical developmental mechanisms. In summary, it has remained unclear whether the transcriptome contains complex shapes (e.g. loops or cavities) during development, and if so, how they are used in different organisms.

Algebraic topology is a branch of mathematics used to describe the properties of topological spaces via group theory (15). In the context of developmental biology, topological spaces can formally represent cell states undergoing differentiation. Using this framework a direct hypothesis is that if we view cell states as points in a topological space, there should be a path connecting a progenitor cell (e.g. a stem cell) and a fully differentiated cell. Furthermore, another topological hypothesis is that an oscillatory circuit would generate a 1-dimensional topological feature in phase space, i.e. transcriptome space. In Fig 1 D we show the phase space of a classic dynamical model of a biological oscillator and its corresponding topological interpretation.

In this work, we leveraged classical and modern tools of algebraic topology. In particular, we used topological invariants – the Betti numbers– to identify the function of gene regulatory networks in developmental transcriptome atlases (Fig 1 B). Specifically, we examined (*i*) if developmental atlases are completely connected and, (*ii*) if they are simply connected. Our analysis revealed that (*i*) is predominantly true under mild considerations, as expected. Surprisingly (*ii*) is not always true – we found instances of 1− dimensional topological signatures corresponding to convergent and cyclic gene expression programs. These results provide strong evidence against the tree hypothesis during development (Fig 1 C).

To uncover the genetic drivers of these topological patterns, we developed a pipeline, TopGen, which calculates the generators of homology using algebra over ℤ_2_. Using TopGen, we identified genes that are transiently expressed along developmental cycles. Our approach of using computational homology to study the topological properties of developmental atlases opens new possibilities for understanding the complex processes of cell development and differentiation. More broadly, our method is well-suited to formally study the topological properties of transcriptome spaces across dynamic processes like disease and aging.

## 2 Results

### 2.1 TopGen enables analyzing gene expression signatures of topological structures in transcriptome spaces

Understanding the topology of a physical system provides valuable qualitative information about its dynamics [(16), (17), (18)]. For example, clusters and loops are signatures of fixed points and oscillations. In this work we conceptualize the transcriptome as a topological space and will use the term *topology* to refer to its homotopy type, which can be quantified using Betti numbers (Methods). The Betti numbers were developed to capture the invariant properties of manifolds under continuous deformations, and unveil qualitative properties of the vector fields permitted in the underlying manifold, including bounds on the number of fixed points (see e.g. Poincaré-Hopf theorem). Conceptually, this becomes significant when considering the notion of a Waddington Lanscape mathematically, i.e. envisioning development as a flow on a manifold (19).

Previous limitations in interrogating gene expression dynamics during development were overcome with the advent of single-cell transcriptomics. However, current computational methods for the analysis of developmental transcriptomic data often rely on *ad hoc* techniques, characterized by strong topological assumptions or reliance on dimensionality reduction tools like tSNE (20) and UMAP (21). Expanding upon previous reports (14), we analyzed topological signatures before and after employing tSNE and UMAP and found that both methods can both create and destroy homology classes (Fig S6). In the context of biology, these results suggest that oscillatory phenomena could be unnoticed using these methods. Moreover, constraints on metabolic or gene regulatory networks might produce higher-dimensional topological signatures. We developed a biocircuit that displays an *H*_2_ signature (Methods), revealing that higher-dimensional homology identification can aid in the discovery of novel biological phenomena (Figure S7). As a whole, these findings emphasize the necessity of new tools to investigate the topology of biological systems.

Inferring the topology of a manifold given a finite sample has been a long-standing mathematical and computational challenge. This challenge is elegantly captured by the results of Niyogi-Smale-Weinberger [(22), (23)], which state that given a manifold *M*, the inference of its homotopy type is not only dependent on data quality (noise and number of samples) but on the geometry around the salient topological features. Intuitively, this result can be understood by imagining that it is easier to “mask out” the “void” of an ellipsoid by adding noise along its minor axis *a*_minor_, than that of a sphere by assuming the radius *r* of the sphere to be equal to the ellipse’s major axis since *a*_minor_ *< a*_major_ = *r*.

Fortunately, the development of persistent homology has enabled the identification of prominent topological features of a manifold from a data sample [(24), (25)]. In essence, the persistent homology algorithm computes Betti numbers at different increasing radii and records when features (homology classes) appear and decay in the persistent diagram. The crucial topological signatures will be the persistent homology classes with a prolonged lifetime, indicating robustness to noise. To compute persistent homology we leverage the efficient implementation of Ripser (26) in python (27), which employs important theoretical tools for efficiency, such as the use of cohomology (28) and discrete Morse theory (29).

To further investigate topological properties in developmental transcriptome spaces we developed a framework that enables explicitly computing topological generators and their associated gene expression patterns (Fig 1 E). First, our approach utilizes persistent homology to identify the homotopy type of a dataset via the computation of its persistent Betti numbers. Secondly, we developed TopGen, a method that uses the representatives of homology groups to analyze gene expression patterns. In essence, the method involves establishing a common basis for the kernel and image of consecutive boundary maps via the Smith Normal Form (Methods). By calculating the *n*− th Betti number, we can determine the homology group generator from this shared basis. To identify transiently expressed genes, we analyze the mutual information of gene expression and the Laplacian eigenvectors of the homology group generator. Genes with high mutual information indicate transient expression along the topological motif. By hypothesis, cyclic topologies would have oscillatory genes that are transiently active in different parts of the cycle. Furthermore, the eigenfunctions of the Laplace-Beltrami operator encode the geometry of a manifold in an orthogonal basis of harmonic functions, which are by definition, oscillatory (30). The discrete version of these harmonic eigenfunctions also turn out to have oscillatory behavior and are eigenvectors of the discrete Laplacian. Therefore transiently expressed genes had high statistical dependence with the eigenvectors of the Laplacian of the homology generator.

### 2.2 Validating TopGen using a ground truth gene regulatory network

To evaluate the efficacy of our approach, we conducted simulations using dyngen, a software package that utilizes the Gillespie algorithm and real data statistics to simulate the acquisition of single-cell RNAseq data with a user-specified gene regulatory program. We designed a GRN consisting of 100 transcription factors, 10 target genes, and 50 housekeeping (HK) genes, and its wiring diagram is visualized in (Fig S1 A). We verified that the dataset had Poisson statistics, characteristic of single-cell data (Fig S1 B).

To determine whether our pipeline could correctly identify the topology of the dataset, we employed persistent homology, a mathematical tool for identifying prominent topological features (Methods). The persistence diagram revealed that 0− homology classes could not be well separated indicative of a large connected component subject to noise. Furthermore, the persistent diagram also showed the presence of a persistent 1− homology class i.e. a loop (Fig 1 E. orange dots). Together, these results demonstrate that persistent homology is a robust method to identify the topological signature of a noisy single-cell transcriptome dataset.

In order to evaluate the statistical robustness of our approach, we developed a permutation test to provide an uncertainty estimate for our results (Methods). In brief, we asked if the topological feature of a test dataset could be explained by chance. To answer this question, we set out to test the null hypothesis that the difference between the lifetime of the maximal *H*_1_ feature of a test dataset and a simulated tree was null, versus the alternative of the maximal *H*_1_ feature being more prominent in the cyclic dataset. Interestingly, we found that the difference between the simulated cyclic data and the tree dataset was significant(P-value *<* 10^− 4^). In the SI we show a systematic evaluation of this approach using both positive and negative controls (Fig S2).

Next, we utilized TopGen to analyze transient gene expression patterns along the identified 1− dimensional homology class. TopGen enabled us to identify transcription factors and target genes exclusively, while retrieving no housekeeping genes as a negative control (Fig S1 D). Our analysis revealed that the expression of HK genes had low mutual information with Laplacian eigenvectors and they were expressed spuriously throughout the loop(Fig S1 D). Furthermore, as a negative control, we computed the persistence diagram on the HK genes only shows no 1 homology classes i.e. no loops (Fig S1 C). As a whole, these results suggest that our analysis pipeline is capable of identifying the correct topological signature of a dataset, elucidating the causes of topological structures using TopGen, while avoiding the retrieval of spurious genes unrelated to the topological signature.

### 2.3 Persistent homology reveals that transcriptome spaces are path-connected but not necessarily simply-connected during development

Based on these findings, we conducted a survey to investigate the topological signatures of the transcriptome across early development across a wide range of eukaryotes (Fig1 B, Table 1). We curated an extensive collection of 11 developmental datasets, including model organisms such as *Danio rerio, Arabidopsis thaliana, Xenopus laevis, Caenorhabditis elegans* and *Mus musculus*. We also incorporated organisms including *Hydra vulgaris, Nematostella vectensis*, and *Schmidtea mediterranea*, for their importance in regenerative medicine. Furthermore, we used the chordate *Ciona savignyi* because of its close evolutionary relationship to vertebrates. The selected datasets are considered golden standards within the field characterized by dense sampling of crucial developmental timepoints. Together, these datasets comprised 864, 640 single-cell gene expression profiles.

**Table 1:** Summary statistics of single-cell developmental atlases and corresponding Betti numbers.

| | Organism | Number of single-cell transcriptomes | $\beta_0$ | $\beta_1$ |
| --- | --- | --- | --- | --- |
| 1 | *A. thaliana* (10) | 107,840 | 1 | 0 |
| 2 | *C. elegans* (2) | 89,701 | 1 | 1 |
| 3 | *C. savignyi* (9) | 767 | 1 | 0 |
| 4 | *D. melanogaster* (7) | 94,315 | 1 | 0 |
| 5 | *D. rerio* (8) | 120,444 | 1 | 0 |
| 6 | *H. vulgaris* (3) | 27,992 | 2 | 2 |
| 7 | *M. musculus* (31) (Weiss) | 92,649 | 1 | 0 |
| 8 | *N. vectensis* (4) | 37,712 | 1 | 2 |
| 9 | *S. mediterranea* (6) | 6,394 | 1 | 0 |
| 10 | *X. laevis* (5) | 188,020 | 1 | 0 |
| 11 | *X. tropicalis* (1) | 98,806 | 1 | 0 |

To interrogate the validity of the tree hypothesis, we conducted persistent homology analysis on our developmental atlas compendium. In order to assess whether the transcriptome space represents a topological tree, we focused on dimensions 0 and 1, as a tree is characterized by Betti numbers *β*_0_ = 1, *β*_*i*_ = 0 for all *i >* 0. Specifically, if the transcriptome space has *β*_0_ *>* 1, this would mean that the topological space is not path-connected and would provide evidence against the tree hypothesis. Furthermore, since trees are defined as acyclic structures, a value of *β*_1_ ≥ 1 would strongly suggest that the underlying topological space is not a tree. To answer the question of path-connectedness in development, we developed a method that leveraged the special nature of the 0− th persistent homology. The 0− th *persistent* homology is special since all persistent 0− homology classes are born at the start of the filtration. Thus, the most parsimonious 0− homology of the data would thus appear as a gap on the ordered lifetimes, and is equal to a maximum on the graph of the second differences of the ordered lifetimes (Methods). We benchmarked this approach by simulating clusters in high-dimensional spaces (SI). Using this approach we found that all but the Hydra atlas had more than one connected component (Fig 2). For the hydra dataset the largest components consisted of the main germ layers, endoderm and ectoderm; this result is unsurprising since the hydra cell atlas was not constructed using a timeseries. Together these results suggest that single-cell transcriptomes are predominantly path-connected during development.

**Figure 2:**
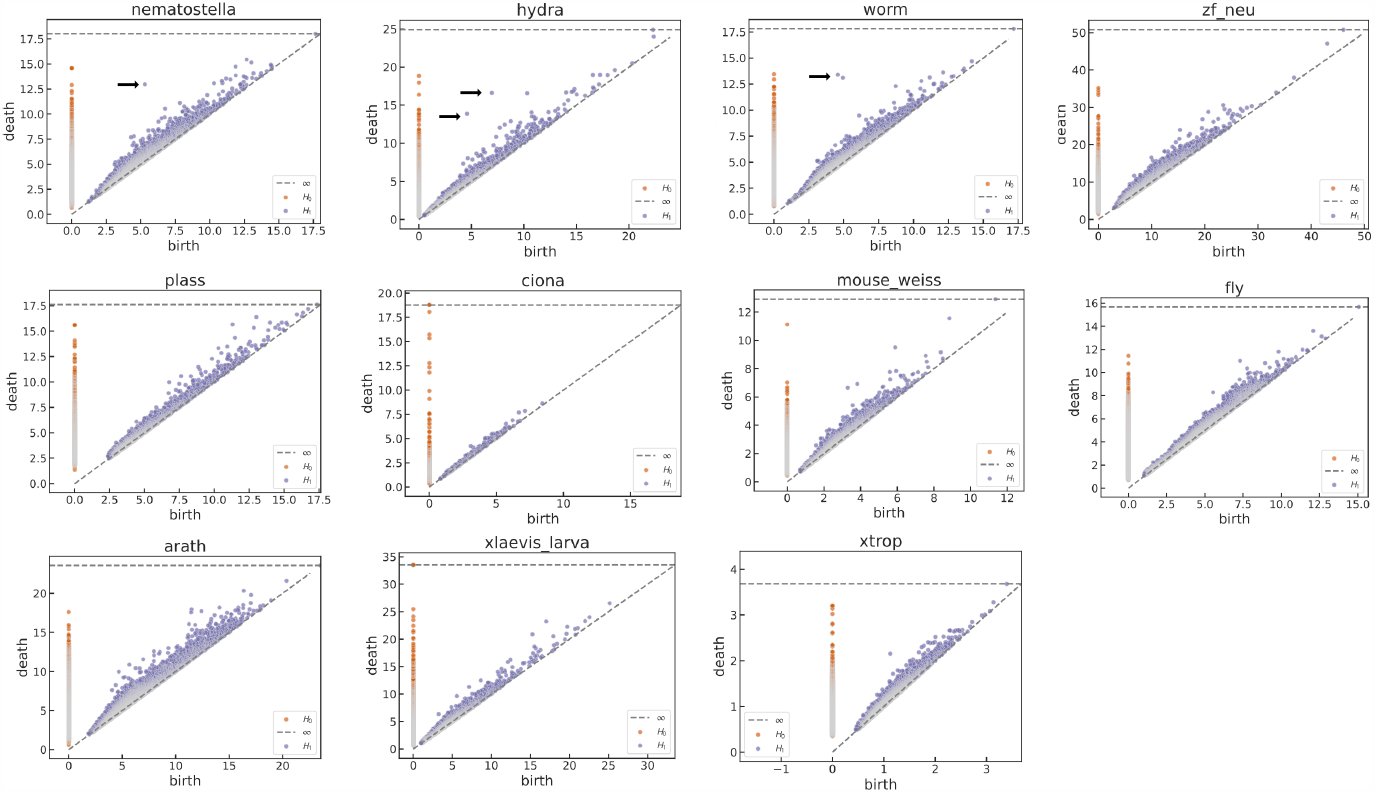
Persistence diagrams for topological single-cell atlas survey. Persistence diagrams summarize the topological features extracted using persistence homology. Each point in a persistence diagram corresponds to a persistent homology class, where its coordinates are the radius at which the feature appeared (*x*-axis) and when it ceased (*y*-axis). Highlighted are *H*_1_ homology classes found in this study.

We continued our topological investigation via the analysis of 1− homology. The alternative to the tree hypothesis is strongly motivated by the 1− homological signature of oscillators in phase space (Fig 1 D). In development, oscillators are implemented e.g. in the somitogenesis circuit. Previous studies have conjectured that the lineage history of single cells could display complex topologies such as cycles or loops (32). Transcriptomic profiling studies have demonstrated that the formation of a gene expression loop in the cell cycle [(33), (34)] when examining solely cell cycle genes. Other studies have used *ad hoc* methods (1) that specify a topology (e.g. a tree) or used dimensionality reduction tools that can strongly distort the data topology. To the best of our knowledge, the unbiased discovery of cyclic topologies has yet to be formally reported in the literature.

To formally, test the tree hypothesis, we used our topological permutation test described in the previous section (Methods). To our surprise, we were able to identify *H*_1_ classes in three datasets (Fig 2): the seam cells of the Worm - representing a stem-like cycle (P-value = 8*×*10^− 3^), in the gland cells of *H*.*vulgaris* (P-value = 0.037) and in the cnidocytes of the cnidarians *H*.*vulgaris* (P-value *<* 10^− 4^) and *N*.*vectensis* (P-value *<* 10^− 4^) which we explain below.

### 2.4 A topological feature could support transdifferentiation of zymogen gland cells to mucous gland cells in Hydra

To begin the exposition of our case studies, in this section we’ll describe evidence for a topological signature providing support for transdifferentiation in the cnidarian *Hydra vulgaris*. Hydra is a 1 cm long freshwater organism that has the extraordinary feature of full-body regeneration. Classic experiments by Campbell in the 60s showed that the hydra can replace the entirety of its cell repertoire ≈ every 20 days (35). Another way to say this is that Hydra has remarkable cellular *stemness*, provided by a particular cell type called interstitial stem cells (ISCs) which can replenish virtually all main cell types of the organism: germ cells, neurons, gland cells, and nematocytes.

Homeostatic self-renewal of Hydra enabled Siebert et al.(3) to construct a cell atlas spanning crucial developmental stages by sampling organisms on different days. In total, they reported more than 27, 000 single-cell transcriptome profiles and were able to sample the main layers of endoderm and ectoderm, as well as low abundance cells such as neurons and stem cells.

Evidence that head mucous cells arose from interstitial stem cells was present in early molecular studies of the hydra (36). ISCs however were reported to be predominantly in the gastric region (low head and foot). Siebert et al. (36) resolved this conundrum by showing that zymogen gland cells(ZMGCs) present in the body could transdifferentiate into granular mucous gland cells (GMGCs). Therefore, the topology of the transdifferentiation mechanism predicts that there would be a corresponding homology class present in transcriptome space (Fig 2b). We applied persistent homology and found persistent homology groups with large lifetimes in this dataset (Fig 2). We found that the most persistent homology was statistically significant as compared to a null tree topology (P-value = 0.038). After thorough analysis, we found that the two persistent homology classes indeed corresponded to gland cells and nematocytes; we explain the latter in the following section. We applied TopGen and found important known marker genes involved in gland cell function and some uncharacterized genes.

We developed a visualization (Methods) to display TopGenes corresponding to a *H*_1_ homology class. Inbrief, we clustered genes and used the first Laplacian eigenfunction as a coordinate for the geometry of the homology class. Finally, we classified genes as early (corresponding to ZMGCs) middle (corresponding to spumous mucous gland cells), and late for granulous mucous gland cells (GMGCs). Note that this classification does not correspond to “pseudotime” but to the geometry of the loop. For instance, TopGen identified multiple digestive enzymes in early and middle gene sets including peptidases (CBPA2, NAS1, CTRC), glycosidases (HEXC), and chitinases (CHIA, CHI13). In contrast, we found multiple mucin homologs (MUC2-RAT, MUC5B-CHICK) contained in “late” gene sets in agreement with the function of GMGCs. Together, these results suggest that TopGen identifies genes crucial to the function of gland cells and that the topology of the transdifferentiation circuit is congruent to the observed data topology in transcriptome space.

### 2.5 A conserved convergent circuit in cnidocytes

As mentioned in the previous section, we found a persistent 1− homology class in the nematocytes of the cnidarian *Hydra vulgaris*. Interestingly, using persistent homology, we found non-trivial persistent homology in another cnidarian, *Nematostella vectensis*. The second case study, which we explain in this section, expands on these discoveries.

Despite their seemingly simple anatomy, cnidarians possess a complex genomic and regulatory repertoire. Cnidocytes are specialized cell types that possess toxins that enable cnidarians to catch prey and contend with predators and thus are the characteristic cell type of cnidarians. From an evolutionary perspective, cnidocytes constitute an essential innovation for the prevalence of this phylum. Furthermore, many cnidarians enjoy remarkable regenerative properties. Together, these features make cnidarians attractive model organisms for studying development.

In an effort to uncover the regulatory mechanisms of cell type diversification across development in *N. vectensis*, a cell atlas was recently performed (4). This Nematostella cell atlas contains samples of early development comprising samples from the gastrula (18, 24 hpf), planula (2,3,4 dpf), and polyp (5,8,16 dpf) stages. Furthermore, they included data from adult tissues of the pharynx, body wall, mesentery, and tentacles. This dataset represents the most exhaustive resource to date for *N. vectensis* single-cell transcriptomic diversity.

We applied our topological analysis approach and found that the *N. vectensis* dataset contains some prominent *H*_1_ persistent features. After some analysis (SI), we found that the cnidocytes have a topological signature of a wedge sum of two circles. We found no H2 persistent feature discarding the possibility of the topological equivalence to a 2D topological manifold like a torus. Interestingly, these topological features were reported in the original study (4) but UMAP increased the number of *H*_1_ features. Consistent with their findings, we found two regions of cnidocytes, one of which did not contain cnidocytes from the gastrula samples (mature). Interestingly, the fraction of mature cnidocytes was higher for the second generator (Fig 4 B, D).

**Figure 3:**
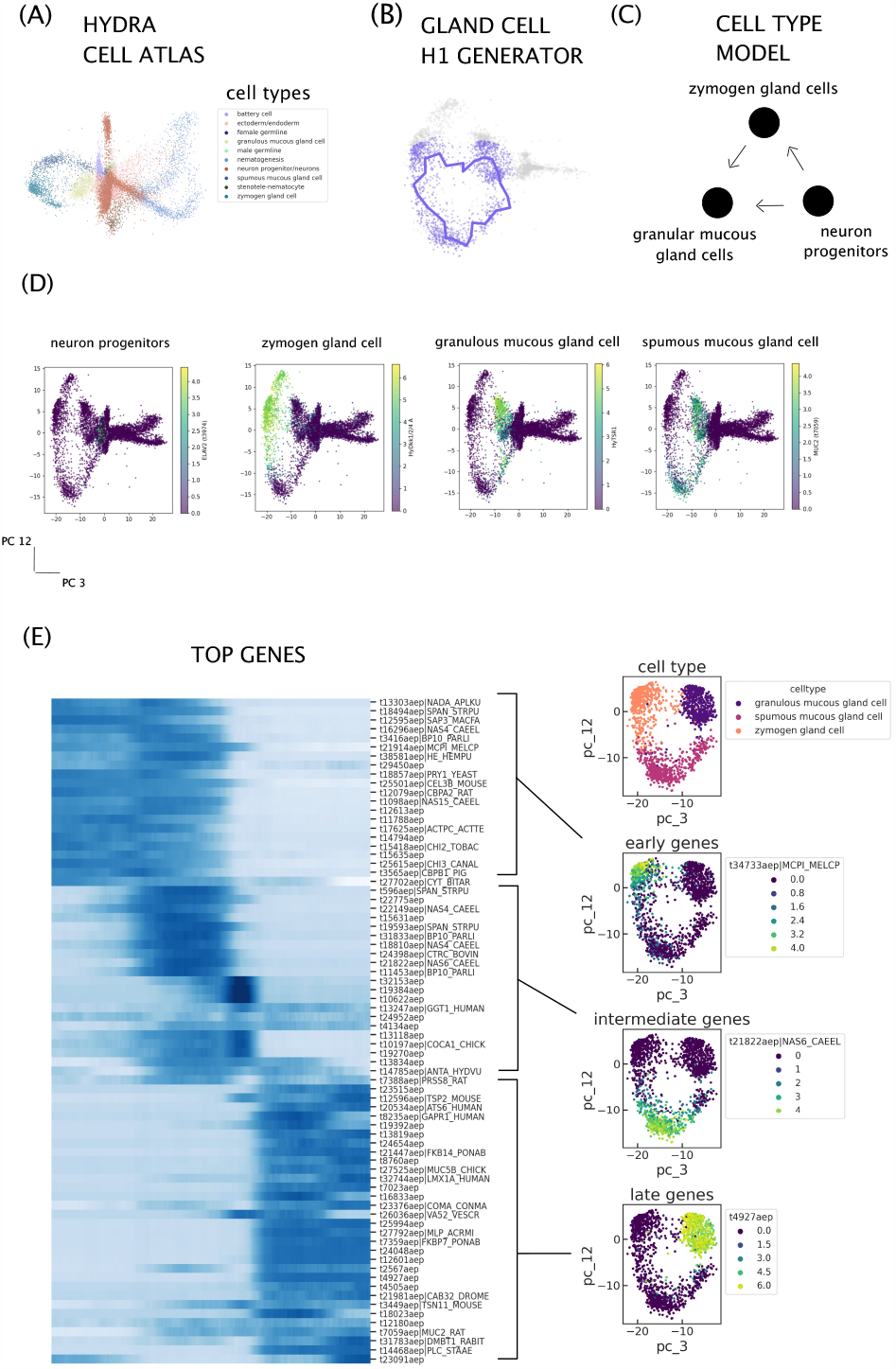
A topological feature could support transdifferentiation of zymogen gland cells to mucous gland cells in *Hydra*. (A) Hydra cell atlas colored by cell types. The plot shows cells projected onto principal components 2 and 13. (B) Inferred homology generator using TopGen. Highlighted are the cells in the neighborhood of the 0− skeleton of the homology class representative. (C) Convergent cell type model as specified by (3). (D) PCA plot for the main gland cell types and neuron progenitors, colored by the expression of their corresponding cell markers (neuron progenitors: *ELAV2*, zymogen gland cells: *HyDkk1/2/4 A*, granular mucous gland cells: *HyTSR1*, spumous mucous gland cells: *MUC2*). (E) TopGenes are ordered by their expression along the homology class (left). Cells in the neighborhood of the homology group representative (right).

**Figure 4:**
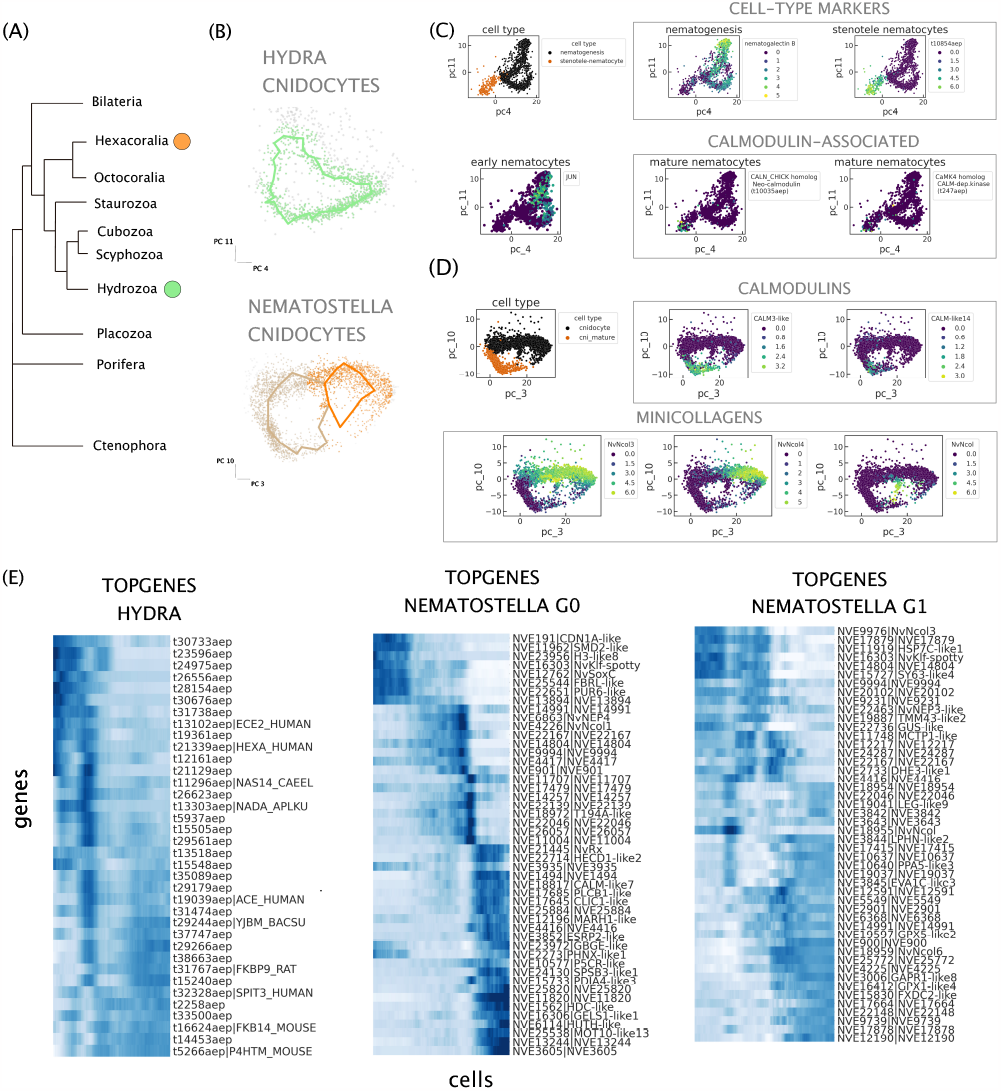
A conserved convergent cycle in the cnidocytes of *H. vulgaris* and *N. vectensis*. (A) Coarse-grained phylogenetic tree diagram of the evolutionary relationship of *Nematostella vectensis* and *Hydra vulgaris*. Both *N. vectensis* and *H. vulgaris* are part of the cnidarian phylum (light orange square), and members of the Hexacoralia and Hydrozoa classes respectively. (B) Homology generators for *Hydra* (top) and *Nematostella* (bottom) using TopGen. (C) Cnidocytes of *Hydra* colored by cell type and developmental markers. PCA plot of cnidocytes colored by markers of stenotele subtype (t10854aep) and during nematogenesis (nematogalectin B) and the corresponding classification (top) by the authors (3). Cnidocytes are colored by the expression of early (JUN transcription factor) and late (calcineurin/calmodulin genes) developmental markers (bottom). (D) Cnidocytes of *Nematostella* colored by calmodulin (top) and minicollagen gene expression (bottom) and their corresponding cell type classification by (4). (E) TopGen enables finding transiently expressed genes along the topological motif.

To investigate gene expression patterns across the topological motifs we found in *Nematostella*, we applied TopGen. Cnidae are the organelles of cnidocytes that contain a tubule ejected upon mechanical input. The cnidocyst tubules are composed of minicollagens, nematogalectins, and other structural proteins that give these macrostructures its functional properties for prey capture and defense. We found that the expression of minicollagen proteins differed across the two topological motifs in cnidocytes, suggesting that this molecular diversity could represent the vast morphological diversity of cnidocytes (Fig 4 D).

In order to understand the regulation of this diversity we asked if there were differences in the transcription factors contained in the TopGenes found by TopGen. Interestingly, we found that the TFs CnidoFos, Pou4, and SoxA in the set of top genes of the first generator, while Jun was in the set of the second generator (Fig S3). This is in agreement with results in bulk measurements where Jun was overexpressed 3-fold while Fos was overexpressed 16-fold than control in mature cnidocytes (37). Furthermore, Pou4 has been reported as a regulator of cnidocyte terminal differentiation (38), where Pou4 mutants produce NvCol3 minicollagen (marker of cnidocytes) but fail to assemble mature cnidocysts. In contrast, Jun knockdowns largely lacked the expression of NvCol3, suggestive of disruption of early nematogenesis.

### 2.6 A stem-like fate maintenance circuit drives an *H*_1_ homology class and enables identification of glial fate priming in early development

The development of the worm *C. elegans* is invariant and has been mapped cell by cell (39), and thus there is a plethora of knowledge of its developmental cell biology. Seam cells are lateral hypodermal cells and perform a stem-like fate. Seam cells display proliferative, symmetric cell division to expand the stem-like pool and asymmetric division to differentiate into hypodermal cells (40). Furthermore, seam cells have an essential function in development by secreting proteins that help the worm elongate and molt. Finally, some seam cells develop into neuron programs for e.g. development of the deirid. The transcription factors ELT-1, RNT-1, and BRO-1 control the ratio of symmetric and asymmetric cell divisions (41). This, tight control of these cell divisions is crucial for properly developing the hypodermis and parts of the sensory system in *C. elegans*.

The C. elegans developmental atlas (2) comprises the first 12 hours of development, representing the worm’s lifetime from the first few cell divisions up until the beginning of the L1 larva stage. Using persistent homology, we identified a prominent *H*_1_ homology class (Fig 2). Furthermore, we identified that the cycle consisted predominantly of seam cells and hypodermal cells (Fig 5 A,D). There was a second persistent homology class identified by PH corresponding to muscle cells, but it turned out that geometrically it was a ruptured circle. The seam cell loop was not reported in the original study and could be destroyed using UMAP projection (Fig S8). Interestingly, this 1-dimensional homology class is not related to the cell cycle as it contains from the early stages of gastrulation (*≈*200 minutes after cleavage) up to the beginning of the L1 stage (12 hours after cleavage). Furthermore, persistent homology analysis of cell cycle genes did not contain 1-dimensional topological signatures (Fig S4).

**Figure 5:**
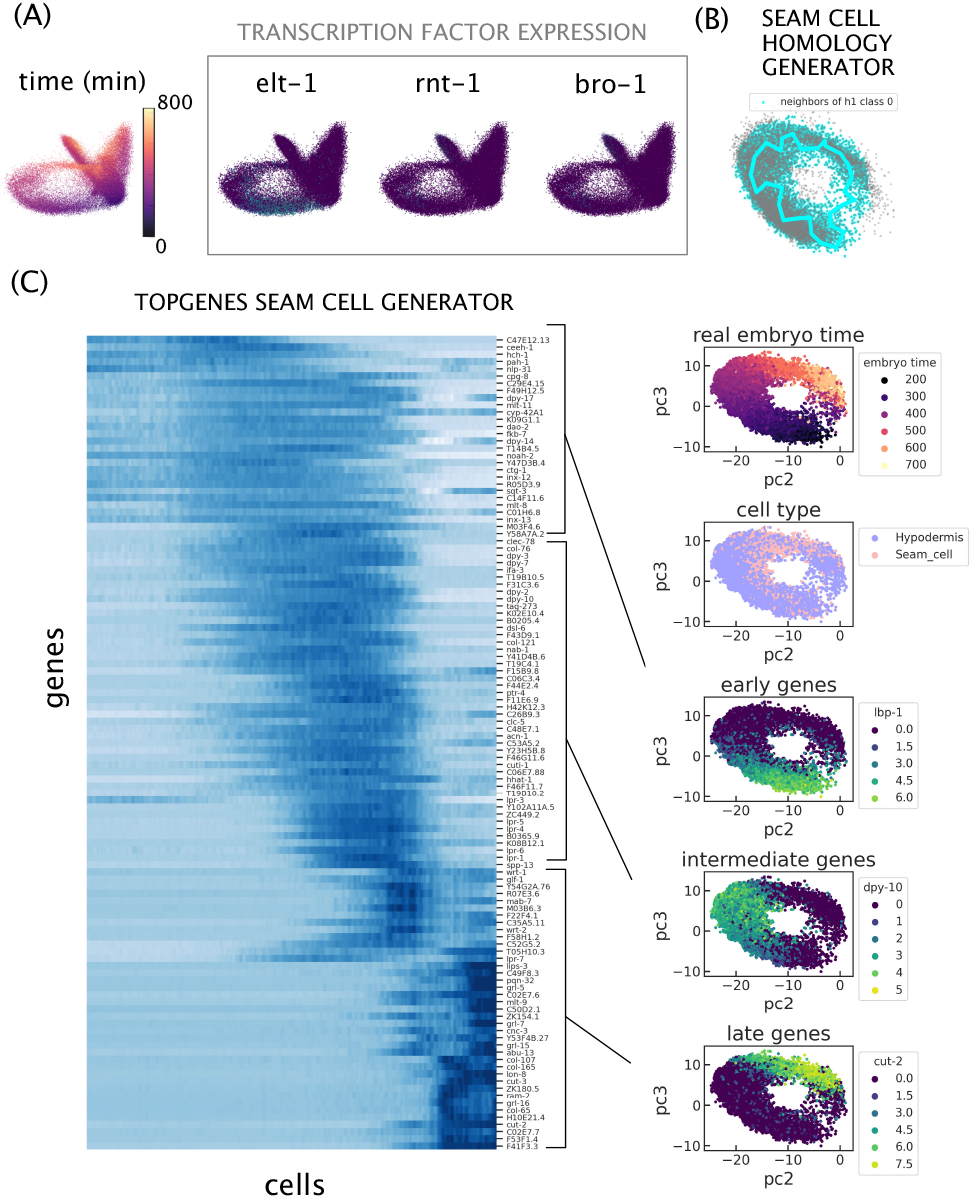
Stemness maintenance is encoded as a cycle in transcriptome space in *C.elegans*’ seam cells. (A) PCA plot of the *C*.*elegans* developmental cell atlas, colored by time after hatching in minutes (left), and by the transcription factors elt-1, rnt-1 and bro-1 respectively. (B) Detected homology class of the seam cells. (C) Top genes ordered by their activation around the *H*_1_ homology class.

We found that consistent with the regulatory events of seam cell regulation (41), the transcription factor ELT-1 is expressed early in the cycle from 100-400 minutes after the first cleavage (Fig 5 A). Moreover, we found that the transcription factors BRO-1 and RNT-1 are expressed in the intermediate part of the cycle from approximately 300-600 minutes after the first cleavage event. Further, glial cells were present at the end of the cycle, which is consistent with the possible cell fates of seam cells. Together, these results support the hypothesis that the *H*_1_ homology class is a topological signature of the seam cell stemness cycle.

We applied TopGen to the *C. elegans* atlas and found fasn-1, cutl-2, and noah-1 to be transiently expressed early in the cycle. Importantly, noah-1 knockdown embryos fail to elongate and rupture (42). We also found genes expressed along the intermediate stages of the homology class such as the genes mlt-8, mlt-9, and mlt-11 which are essential for molting (43). Finally, we also discovered genes expressed in all but the initial and terminal regions of the cycle, such as sqt-3, which has been reported to be essential for locomotion and viability (44). As a whole, these results suggest that the homology class is driven by both structural and mechanical functions of seam cells. These functions are essential for the worm prior to hatching.

## 3 Disussion

In this work we viewed scRNAseq landscapes as topological spaces. Furthermore, we discovered and quantified that cell transcriptome spaces are not always simply connected, i.e. can have 1-dimensional holes. This fundamentally novel concept challenges our prior understanding of development as a purely branching process.

In the broader context beyond development, previous studies have identified topological loops. However, these studies relied on subsetting cell cycle genes [(33), (34)] to identify loops, or leaned heavily on the processing of the data. These previous efforts have not yet described the employment of tools capable of rigorously discovering topological loops in cell state manifolds, let alone to explore gene expression in these non-trivial topological structures. Therefore our approach represents an unbiased way to discover and dissect gene expression in topological features from high-dimensional transcriptome data, which had not yet been achieved by previous studies. Moreover, it is probable that some of the initial studies, which provided the data for our analysis, may not have identified topological motifs due to their reliance on UMAP and tSNE in their analysis pipelines, potentially introducing topological distortions (Fig S8).

One remarkable result from our analyses is that loops are rare occurrences, present in only a small fraction of our analyzed datasets. As our resolution to probe the transcriptome increases, we predict that more topological signatures will be uncovered across different organisms. This underscores the need for further research to develop experimental and computational techniques to expand upon these findings.

The potential that a disconnected manifold could become connected noise coud alter the topology, remains a possibility, however, this scenario is challenging to discern without a parsimonious null hypothesis. For instance, in the context of development, the parsimonious hypothesis for zero homology is that the transcriptome is path connected. In contrast, in other contexts, it may be easier to use alternative hypotheses, such as in fully differentiated systems, timecourses of disease progression, perturbations, and aging.

A concrete example of the possibility of noise phenomena affecting the topology in our analyses is the case of cnidocytes in Nematostella. This is particularly subtle, since the NSW theorem predicts that the manifold is conditioned by the minimal distance to its medial axis, which in this case is a principal component of a small singular value (PC 13). Geometrically this can be seen by noting that the topology of the figure eight is formed by drawing a path across the minor axis (PC13) of an ellipse. However, functionally, we showed that gene expression patterns are fundamentally different across the two 1-homology generators which strongly suggests that this topology is prominent. These findings open up new avenues for future research incorporating the analysis of noise and topology. In particular, using a dynamical model of the system with a topological correspondence could help ellaborate more complex hypothesis testing regarding the noise in the system.

Finally, our approaches for finding transiently expressed genes could be modified to yield different results. There are other approaches that could exploit the experimental sampling time or spatial information to discover gene expression patterns (45). Other class of models could incorporate different methods of gene analysis. For example, Siebert et al. (3) used TF motif search to retrieve potential regulators for cell fate. This analysis in the Hydra yielded an enrichment of Pax2a and RFX regulators in gland cells and cnidocytes respectively. In contrast, we found JUN to be transiently expressed in gland cells and LMX in cnidocytes. Thus our analysis provides potentially complementary information about complex biological processes as compared with other analysis methods.

Limitations of our study include the incomplete acquisition of all cell circuits in the analyzed organisms by the limited resolution of scRNAseq: low mRNA capture rate, biochemical noise, measurement error, and low sequencing depth all affect the identification of homology. Another limitation is that we limit our scope to the study of 0 and 1 homology, because of the high computational cost for higher dimensions. For instance, the calculation of 2− homology requires the computation of the boundary map *∂*_3_ mapping 3−simplices to 2− simplices; for *n* =10^5^ datapoints (as is the case for some of our atlases) the number of possible 3− simplices to is on the order of 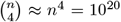, at the end of the filtration. Thus, even with the efficient implementations to date, memory requirements become prohibitively large for higher dimensional homology. It is important to underscore that higher homology could encode important biological phenomena that still awaits our theoretical understanding. In the SI we show how composition of a simple oscillatory circuit with variable repressor strength can lead to 2− homology. Our approach generalizes for higher dimensional homology and we thus envision that future research in computational algebraic topology will enable the computation of higher dimensional homology for single cell developmental atlases.

In conclusion, this work has shed light on the potential of viewing scRNAseq landscapes as topological spaces, offering a new way to understand complex biological processes and uncover hidden patterns. Beyond its immediate impact, our approach holds promise in complementing traditional bioinformatic analyses, opening up new avenues for exploring complex biological phenomena like aging and disease. For instance, the discovery of the stemness loop in *C. elegans* provides a new way to understand the mechanisms of pluripotency maintenance in stem-like cells. Moreover, the implications of our findings extend to cell therapy, where a deeper understanding of the topological signatures of stem-like properties may facilitate engineering strategies using synthetic biology. In the context of cancer, cycling stem cells may be unaffected by approaches targetting fast-growing cells. Because they may be relatively protected from current treatment strategies, cancer stem cells are thought to be responsible for resistance to chemotherapy and the recurrence of disease. By harnessing the power of topology in scRNAseq analysis, we have set a trajectory for advancing our understanding of complex biological systems and paving the way for innovative therapeutic strategies.

## 4 Methods

### 4.1 Topological model of transcriptome data

Let *X* = {*x*_1_, …, *x*_*m*_, *x*_*i*_}*∈* ℝ^*n*^ be a set of *m* transcriptome profiles where *n* is the number of genes. We consider the set of transcriptomes *X* to be points sampled from an underlying manifold, *M*, via a measurement process that generates data points *x* drawn probabilistically through a measurement process *P* (*x*|*y*) where *x∈M* and *y∈T*_*x*_*M* ^*⊥*^ (23). Our goal is to infer the underlying topology of this manifold through analysis of the sampled points. We are interested in the topology of the manifold because topological structure can reveal principles of gene regulation and cell-state control. Formally, our goal is to estimate the homology groups and corresponding Betti numbers of the underlying manifold *M*. The Betti numbers encode, informally, the number of holes of increasing dimension.

Intuitively, each point in *X* can be viewed as a representative of a small neighborhood of the transcriptional manifold *M*. By forming open sets around representative points, our aim was to cover the manifold by including nearby transcriptional states. In other words, an open cover of *X* would represent oversampled but continuous transitions of cell-states nearby the sampled single-cell transcriptomes. This approach would in principle allow us to assess the local structure of the manifold *M* using the sampled data *X* and infer of its underlying topology.

Fortunately, the topology of *M* is encoded in the simplicial complex built from the intersection signature of an open cover of *X* (SI) (15). We can thus infer the topology of *M* by associating a simplicial complex structure to *X*. In brief, a simplicial complex is the discretization of a manifold. For instance, 2D manifold (a surface) can be triangulated, effectively resulting in a simplicial complex. The power of simplicial complexes lies in their computational capabilities, allowing for straightforward computation of homology. The building blocks of simplicial complexes are simplices, which are generalizations of triangles: 0 − simplices are points, 1− simplices are line segments, 2− simplices are triangles, 3− simplices are tetrahedrons, and in general, *p*− simplices are *p*− dimensional polytopes. In our framework, a transcriptional 2− simplex would be a set of three nearby transcriptomes that can continuously deformed into each other, and that locally represent a 2D neighborhood of *M*.

It turns out that for computational purposes, it suffices to use only the indices of the transcriptomes forming the simplices, forming an abstract simplicial complex (ASC). Formally, an abstract simplicial complex *K* is just a collection of sets that is closed under the action of subsetting, i.e. if we let *σ*= {*x*_0_, *x*_1_, …, *x*_*p*_ } be a *p*− simplex of *K* and *τ* ⊆ *σ*then *τ* is in *K* automatically. The most efficient procedure to build an ASC from data is the Vietoris-Rips (VR) algorithm. This algorithm was first developed to define a homology theory for arbitrary metric spaces. Thus, to effectively infer the topology of the transcriptional manifold *M* using the VR algorithm, we just need our transcriptome data *X* and a distance metric *d*(,). For all experiments in this work we used the euclidean metric. The VR complex VR(*X, ε*) is constructed as follows:

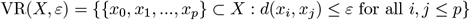

i.e. *V R*(*X, ε*) is a family of subsets of *X*, abstract *p*− simplices, defined if all pairwise distances of *p* + 1 vertices are less than or equal to *ε*.

In summary, abstract simplicial complexes enabled the calculation of homology groups via simplicial homology theory. The core of the homology calculations lie in the definition of a *chain complex* which we explain in the following section.

### 4.2 Algebraic topology framework to compute homology

A chain complex is an algebraic structure that enables defining and calculating homology groups from simplicial complexes. Formally, in the context of simplicial homology, a chain complex is a sequence *{*(*C*_*p*_, *∂*_*p*_)*}*_*p*=0,1,…,*n*_, where *C*_*p*_ are abelian groups generated by linear integer combinations of *p*− simplices, and corresponding group homomorphisms *∂*_*p*_, the boundary maps, which encode how *p*− simplices are connected to (*p* − 1)− simplices (SI). In other words, the chain complex contains all the information to construct the simplicial complex by “gluing” its building blocks. We denote the chain complex by:

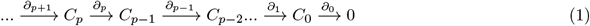

The most important property of this algebraic structure is that *∂*_*p*− 1_ *∂*_*p*_ = 0, i.e. the composition of two consecutive boundary maps is equal to the zero map. This formalizes the geometric concept that taking the boundary of a manifold (with boundary) yields a submanifold that is boundaryless. This immediately implies that im*∂*_*p*+1_ *⊂* ker*∂*_*p*_.

Precisely this property allows us to define the *n*− th homology group *H*_*n*_ and the corresponding *n*− th Betti number as:

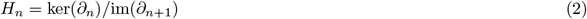

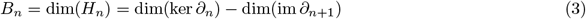

The geometric intuition of this property is that the boundary of a manifold without boundary has no boundary. For example, the 1− dimensional homology will be equal to the number of cycles not coming from the boundaries of triangles. To unravel this argument please note that 1− simplices are edges and 1− dimensional cycles are combinations of edges that return to the node of origin, will be in the kernel of *∂*_1_. Furthermore, edges of triangles present in the simplicial complex would be in the image of *∂*_2_. Thus subtracting the number of triangles (dim im *∂*_2_) from the number of cycles (dim ker *∂*_1_) will thus give us the number of 1− dimensional holes or loops. This idea generalizes for high dimensional holes.

### 4.3 Identifying the topological signature of noisy data using persistent homology

To identify the topological signatures from our transcriptome datasets, we used persistent homology (PH) (25). Persistence homology is a powerful tool for inferring the topology of a dataset that is robust to small noise perturbations. In contrast, classical homology groups can be highly sensitive to noise. The idea of PH is to build a family simplicial complexes by scanning across increasing radii in the VR complex, from which persistent Betti numbers are computed. The Betti numbers that persist will be the essential topological features from the data. We construct the *filtration*:

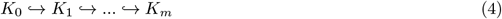

where 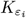 is the VR simplicial complex formed from the set *X* at scale *ε*_*i*_, where for notational convenience we let 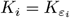. Applying the homology functor to the above sequence, yields a sequence of homology groups (one for each dimension *n*), its **persistence module**, connected by group homomorphisms which are pushforwards of inclusion maps:

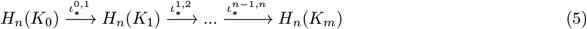

Where the asterisk denotes that *ι*^*\**^ is a function between homology groups. The sequence of homology groups contains the topological features present at different radii. Note that if [*γ*] *∈ H*_*n*_ (*K*_*i*_) then 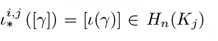, by definition of the induced maps on homology (SI).

The images of the pushforward inclusion maps are the **persistent homology groups**, that is:

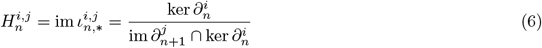

The corresponding *n*− th **persistent Betti numbers** are their corresponding ranks:

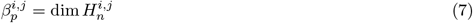

In words, persistent homology groups consist of the homology classes of *K*_*i*_ still alive at *K*_*j*_, and their *lifetime*, or persistence is exactly *j* − *i*. Intuitively, as we increase the scale of the VR complex, holes can be lost (for example when isolated points connect to each other *β*_0_ decreases) or gained (e.g. as edges connect to form a loop *β*_1_ increases). In this way we can consider the image of the inclusion map to analyze the lifetime of homological structures and their associated Betti numbers.

The term lifetime comes from the fact that we can characterize homology classes by when they appear and cease (which is exactly what we’re interested in) : Let [*γ*] *∈ H*_*n*_(*K*_*i*_), we say that the homology class is *born* at *K*_*i*_ if 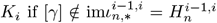. Futhermore a homology class *dies* at *K*_*j*_ if it merges with a previously born homology class exactly at *K*_*j*_, i.e. 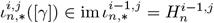 and 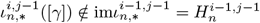.

It turns out that the decomposition of the persistence module in terms if their births and deaths, i.e. its *persistence diagram*, is unique up to isomorphism and completely characterizes the topology of the underlying space. Practically, the persistence diagram can be represented in the plane as pairs (*b, d*)*∈* ℝ^2+^, where *b, d* are the births and deaths of each homology class. Persistent classes far from the diagonal will correspond to those with long lifetimes, and are the classes with strong topological signature. We refer the reader to the review (46) for an in-depth picture on the theory behind persistence homology.

We used the efficient implementation of Ripser in python, pyRipser, to perform our calculations.

#### 4.3.1 Computational scheme to build the chain complex

We use a bottom-up approach to construct the chain complex (*C*_*p*_, *∂*_*p*_) for a Vietoris Rips complex VR(*X, ε*). Homology modulo 2, i.e. homology using algebra over ℤ_2_ is employed for our calculations. The coefficients for *p*− chains represent presence or absence of specific simplices. Our methodology is a modification of the method described in (47).

To initiate the construction, let a distance matrix *D* of size *n×n*, that corresponds to the metric space (*X, d*), where *n* is the number of points in *X*. To obtain the 1− *skeleton*, (i.e. the set o all 1− *simplices* we construct an adjacency matrix using *Ã* = *D <* 2*ε*. The matrix *Ã* will be an *n×n* symmetric binary matrix corresponding to the underlying undirected graph.

To obtain *C*_1_ from *A*, we first remove its redundancy by computing its upper triangle *A* = triu(*A*). We then get the list of 1− simplices by setting *A* to COO sparse matrix format, concatenating the nonzero rows and columns. More generally we define the *p*− simplex matrix as follows: Let the **simplex matrix** *S*_*p*_ be a (dim*C*_*p*_, *p* + 1) matrix that contains the indices of each *p*− simplex in each of its rows.

*S*_*p*_ serves a representation of *C*_*p*_. The boundary matrix *∂*_*p*_ is computed from *S*_*p*_ by computing all 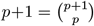 possible (*p* − 1) simplices one can form for each *p*− simplex and store each (*p* − 1)− simplex simplex in a binary matrix precisely representing *∂*_*p*_. In other words, for any given dimension *p* we use *S*_*p*_ to record in each column of the boundary matrix *∂*_*p*_ the set of (*p* − 1)− simplices that each *p*− simplex gives rise to.

For *p ≥* 2, to verify the existence of each *p*− simplex under the VR condition, we need to check that all its (*p* − 1)− simplices are connected. To do this we define an auxiliary matrix *F*_*p*_ as follows:

**Definition. Face** A simplex *τ* is a *face* of *σ*if *τ ⊂ σ*. We say that *σ*is a *coface* of *τ*.

**Definition. Face matrix** *F*_*p*_ The *p*− face matrix *F*_*p*_ is defined as :

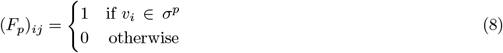

Hence the name *face* matrix: a vertex is naturally a face of a *p*− simplex if it is contained in the simplex.

Now for concreteness, consider the case when *p* = 2 (other cases are an easy extrapolation). Assume a vertex *j* contains more than two incoming edges. For such a vertex, there will be more than two nonzero entries in its corresponding column *A*_*j*_ in *A*. To know if that vertex *j* is contained in a 2− simplex it suffices to check if there is an edge between any pair of incoming vertices. We can now view 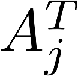 as a *functional* from *C*_0_ *→* ℤ. Thus, given a column of *F*_1_ as input, 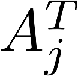 returns the number of incoming vertices into *v*_*j*_ contained in the edge. Thus we can define the map *A*^*T*^ *°F*_*p*_ : *C*_1_ *→* ℤ that has the following property:

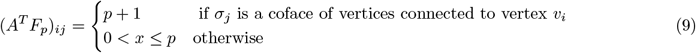

To reiterate, the value of entry (*i, j*) of *A*^*T*^ *F*_*p*_ is equal to *p* + 1 if we form a *p*− simplex of the Rips complex by concatenating the vertices composing simplex *σ*_*i*_ to vertex *j*. Checking where the condition holds gives a simple algorithm to construct *S*_*p*_ for *p ≥* 2. Note that *C*_*p*_ is generated by the simplices stored in *S*_*p*_. Hence, we can construct (*C*_*p*_, *∂*_*p*_) for any dimension as desired.

### 4.4 Calculating homology group representatives

To compute homology groups and Betti numbers we use the Smith Normal Form (SNF) decomposition of each boundary map *∂*_*p*_ over ℤ_2_. More precisely, if we let *∂*_*p*_ be an *m×n* integer matrix, the SNF is a unique factorization 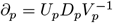 with the following properties:

- *D* is an *m × n* diagonal matrix with ones in the diagonal.
- *U ∈ GL*(*m*, ℤ), *V ∈ GL*(*n*, ℤ), i.e. *U* and *V* are unimodular (with det = *±*1), invertible, integer matrices.
- The number *r* of ones in the diagonal of *D* will be the dimension of im*∂*_*p*_.
- The last *n* − *r* columns of *V* constitute a basis for ker*∂*_*p*_.

The proof of these properties is provided in the appendix. With the above result, we can now easily calculate the Betti numbers. However, since we’re interested in the representatives of homology groups, we need to get the chain groups in the same bases.

Let *D* be the SNF of a matrix *A*. Then define *D*^*\**^ to be the matrix which is the result of permuting the columns of the matrix *D* so that the diagonal block is in the upper right corner, i.e. of the form :

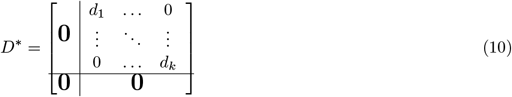

In this sense, we’re thinking of 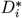 to be the result of applying a change of basis to the boundary matrix *∂*_*i*_ to get 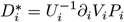.

Accordingly, since operations on columns on *∂*_*i*_, as a change of basis operation, correspond to operations on rows on *∂*_*i*+1_, we have the following definition: we say that 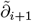 is the matrix into which *∂*_*i*+1_ is carried after applying the operations to diagonalize *∂*_*i*_ using SNF. That is 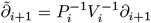. Note that *P*_*i*_, *V*_*i*_ act on the rows of *∂*_*i*+1_.

In a more succint description, let 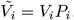, then we define:

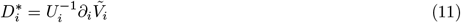

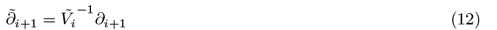

It’s not hard to show that the last *r*_*i*_ rows of 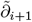consist of zeros, since Smith factor matrices and permutation matrices are isomorphisms we have that 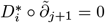(SI). The importance of this result is that by changing bases on a boundary matrix *i* to those specified by the previous matrix following SNF decomposition, we actually advance the SNF of the (*i* + 1)− th matrix and work with the same basis, and come closer to computing the representative of homology.

To do this assume that the SNF of *∂*_*i*_ is available in the form 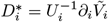.

Change basis to map 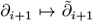, and decompose the matrix using SNF to get 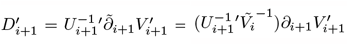

We then have that 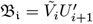 is a common basis for the ker*∂*_*i*_ and the im*∂*_*i*+1_. In particular the columns with index {(dim im*∂*_*i*+1_) + 1, (dim im*∂*_*i*+1_) + 2, …, dim ker*∂*_*i*_} of 𝔅_i_ constitute representatives of the *i*− th homology group (48).

We thus have the following algorithm to compute homology groups:

#### Algorithm 1

Homology group using SNF

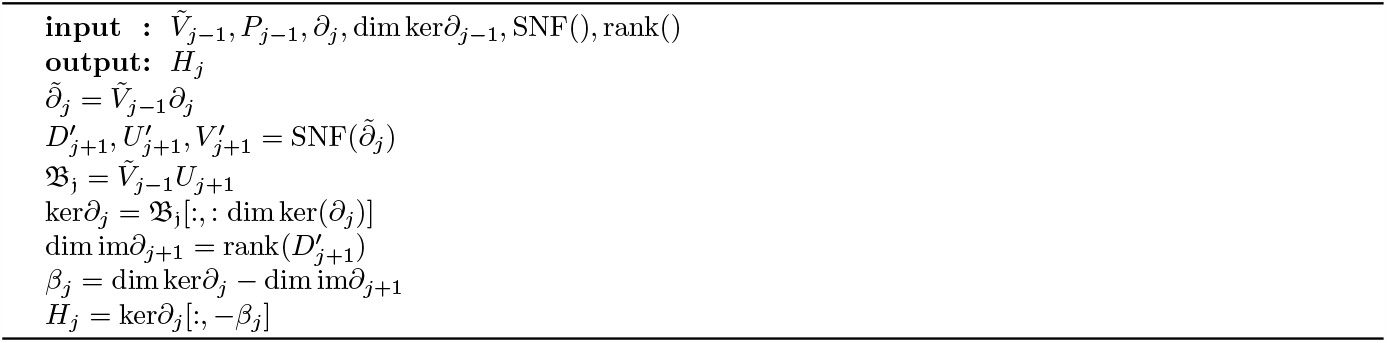

### 4.5 Using homology generators to investigate gene expression

We exploited the homology generators to identify transiently expressed genes along complex topological features. Our methodology is based on the property that Laplacian eigenvectors encode geometrical properties of a manifold (30). Furthermore, eigenvectors can be interpreted as vibration modes, with increasing eigenvalue corresponding to an increasing spatial frequency. We thus extracted transient genes by asking which genes had the highest mutual information w.r.t. the first nonzero eigenvectors of the Laplacian.

### 4.6 Computing Laplacian eigenvectors efficiently using the power method

The Laplacian is a linear operator of dimensions (*p*− simplices, *p*− simplices). For example, *L*_0_ is of size (*n, n*), where *n* is the number of 0− simplices in the simplicial complex. For our puroposes, we just neeeded a small number of eigenvectors from the Laplacian. We thus developed a method to compute the Laplacian eigenvectors using an approximation from the corresponding Markov matrix. Namely, we exploited the fact that spectrum of the Laplacian and the Markov matrix coincide up to a rotation in ℂ (SI), to extract the first eigenvectors of the Laplacian from the eigenvectors corresponding to the largest eigenvalues of the Markov matrix.

### 4.7 Single-cell RNAseq data pre-processing

Single-cell RNA seq count matrices were pre-processed using a standard pipeline. First, we filtered out cells with less than 500 detected genes and 1000 UMIs. To minimize variability in the total number of reads per cell the expression values were normalized using the equation

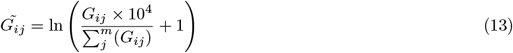

in order to get normalized counts roughly equivalent to those expected in a single-cell.

In addition, we identified the highly variable genes using the coefficient of variation method. Finally, we reduced the dimensionality of the transcriptome by projecting the transcriptome to the first 100 principal components facilitating efficient computation.

### 4.8 Second finite difference method to identify persistent 0− homology

A parsimonious way to retrieve the most prominent 0− homology of a dataset is to select the number of homology features that persist the longest, and that are well separated from the rest of the persistent homology features. In this sense, we would retrieve connected components that are well separated and that have a much larger scale than their possible subclusters. The 0− th homology persistence diagram has the special characteristic that all homology features are born at the start of the filtration. Thus the lifetime equals to the death time for all persistent 0− homology features. This induces a natural order between the 0− homology features, and if we order the lifetimes in descending order, we can naturally form a monotonically decreasing sequence. In other words we would have a sequence {*l*_*i*_}_*i*=0,…,*n*_, where *l*_*i*_ denotes the lifetime of the *i*− th persistent homology group, and furthermore *l*_*i*_ *> l*_*i*+1_. If we let *d*_*i*_ be the difference between consecutive ordered lifetimes, we would have that *d*_*i*_ =*l*_*i*_− *l*_*i*− 1_≤0. We can thus form a sequence of differences {*d*_*i*_ } _*i*=0,…,*n* − 1_, all negative. Therefore, we can get the desired number of prominent persistent homology features by checking when does the difference between consecutive *d*_*i*_’s, i.e. *dd*_*i*_ = *d*_*i*_ − *d*_*i*− 1_. We thus define the parsimonious 0− homology as :

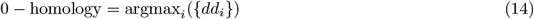

which is equivalent to answering the question: what is the index of the largest gap between consecutive lifetimes of 0− homology features.

### 4.9 Bootstrap permutation test to quantify significance of topological signatures

We used a bootstrap permutation test using a bifurcating tree as a null topology to quantify if the persistent homology signatures could be explained by random chance. To do this we employed difference of maximal lifetime between a test dataset and a null hypothesis dataset. Let the lifetime of a persistent homology class be 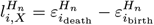 where *i* is the index of the homology class in the filtration.

Our statistical test is based on the following hypotheses:

1. *H*_0_: 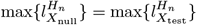
2. *H*_1_: 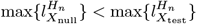

We thus define our test statistic as follows :

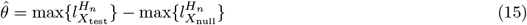

To simulate the null hypothesis, we concatenate the null and test datasets, shuffle, partition, and compute the test statistic 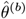 for each bootstrap replicate. We performed *B* = 10^4^ bootstrap replicates of this test and report the P-value as the fraction of simulations in which 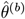 is more extreme than the test statistic 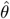. In order for the datasets to be in the same scale, we use the singular values of the test data to scale the principal components of the null dataset prior to all computations.

Additionally, we employed the same test by replacing the maximum, considering the *k-*th order statistic. This enabled generalizing our test for the case when there was more than one persistent feature.

## 5 Supplementary Notes

### 5.1 Benchmarks of topological statistic tests using synthetic data

In order to evaluate the effectiveness of our topological analysis, we conducted simulations of scRNAseq datasets incorporating predetermined data topologies and subjecting them to our topological statistical test. For the purpose of our simulation, we used dyngen, a method that uses the Gillespie algorithm and real data statistics (such as capture rates and library sizes) to mimic the acquisition process of scRNAseq data.

To establish a baseline, we a null hypothesis dataset with a simple bifurcation tree topology. To assess the performance of our method, we performed a positive control experiment with the cyclic gene regulatory topology, and found a significant difference compared to the max H1 lifetimes of the control bifurcation dataset (P-value *<* 10^− 4^). We also performed negative control experiments featuring trifurcation, linear trajectory and binary tree topologies. Our statistical test revealed no significant differences in these datasets: trifurcation P-value = 0.4, linear trajectory P-value = 0.21, binary tree P-value = 0.54 (Fig S1).

Finally, to verify that the test had low false discovery rate, we asked if the second most prominent H1 feature of the the dyngen cyclic topology was significant. For this case, we found that a P-value = 0.24, indicating that the test identifies a single significant topological feature as expected.

For all our experiments, used 10^4^ permutation replicates and 100 principal components for all of our experiments. These parameters were consistently applied across all experiments to ensure consistency and reliability of our results.

### 5.2 Why does tSNE and UMAP break topology?

In this section we give an explanation as to why tSNE and UMAP *can* break topology. For more comprehensive studies analyzing these algorithms we please refer to [(49), (50)]. Both tSNE and UMAP are algorithms for solving an optimization problem. It turns out a term in objective function, in both cases, optimizes for breaking the topology. Let’s begin by using some notation: let *X* ∈ 𝕄(*m, n*) be an *m ×n* matrix of data, where each point **x** ∈ ℝ^*n*^ belong to metric space (*X, d*), and that *d* is the euclidean distance. In general, dimensionality reduction methods aim to find a matrix *Z* ∈ 𝕄(*m, d*) with *nice* properties (where *d << n*). The approach for both methods is to use a proxy of the measure of points being close in both the high and low dimensional spaces, and to minimize the distance between the measures. In both cases, up to affine transformations, the probability that two points in the high-dimensional space are close together will be denoted as *p*_*ij*_, and *q*_*ij*_ for the low-dimensional space. Both methods use modifications of the classical Gaussian affinity :

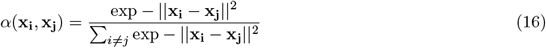

And let the affinities in the high and low dimensional spaces respectively be :

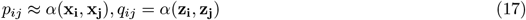

tSNE aims to minimize the Kullback-Leibler divergence between *p* and *q*:

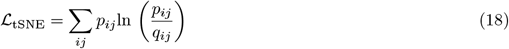

where the optimization can be done using gradient descent using 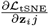. Let’s expand the loss function:

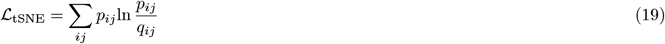

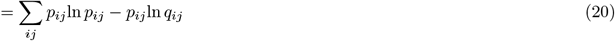

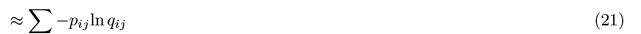

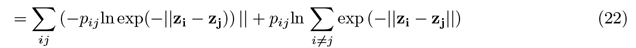

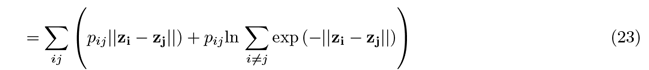

in the third line the approximation is valid since when computing the gradient the term *p*_*ij*_ln *p*_*ij*_ will be eliminated since it only depends on the original data *X*, and not on the low-dimensional representation.

From the last formula we can see that the second will be minimized when ||**z**_**i**_− **z**_**j**_|| ^2^ is large, and hence is called the “repulsive” term in the literature (49). The repulsive term explains the lack of topological preservation of tSNE. In contrast, the first term, also called the “attractive” term, actually is topology-preserving. To understand why the attractive term doesn’t modify the inherent topology of the low-dimensional representation, one has to note that the first term corresponds to the Laplacian embedding: its solution is exactly equal to the smallest nonzero eigenvectors of the Laplacian. It is well known that the graph Laplacian encodes the zero-dimensional homology: the dimension of the kernel is equal to the number of components of the graph. It turns out that the first graph Laplacian eigenvectors also encode the global geometry of the data (30). For a theoretical guarantee we have to look at the higher-order combinatorial Laplacians (23):

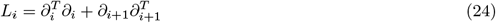

Note that in this context the graph Laplacian will be *L*_0_. The kernel of the *n*− th combinatorial Laplacian will be exactly the *n*− th homology group (23).

On the other hand, UMAP has the following objective function :

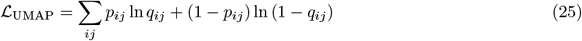

Note that the first term of the above objective function is exactly equal to (22). The second term will also have attractive and repulsive terms by symmetry. Thus UMAP has the same topological problem as tSNE. Furthermore, when using the Laplacian embedding as initialization, UMAP will satisfy the attractive terms exactly and will thus only modify the low-dimensional representation by repulsive forces.

### 5.3 Experiments of topological distortion of dimensionality reduction methods

After verifying that there is no strong topological-preservation guarantee mathematically, we set out to test the effect of the dimensionality reduction methods tSNE and UMAP on topology experimentally. Our effort was to show the worst-case scenario of both algorithms to prevent the misinterpretation of the results of these algorithms. To do this, we decided to use the 1 and 2 spheres in order to visualize the results. The experiments were designed to investigate the behaviour of the algorithms as a map between topological spaces *f* : *X→Y*. Specifically, we wanted to verify if the algorithms could preserve the topology. We reasoned that in real datasets we would not find these platonic manifolds, and thus we decided to apply a homeomorphism prior to applying the algorithms.

For the circle we applied the following homeomorphism:

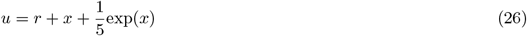

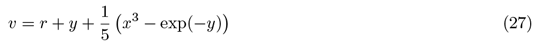

and for the sphere:

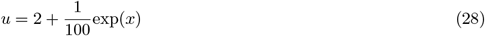

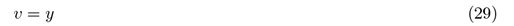

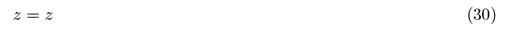

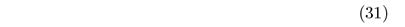

To verify that the above functions are homeomorphisms one can easily check that they are diffeomorphisms by noting that they are differentiable and that their Jacobians are nowhere singular.

Furthermore, previous studies have shown that the perplexity parameter is the strongest control parameter in tSNE (14) for its capacity to preserve or modify the topology of data. We thus decided to vary the perplexity parameter for tSNE and the number of neighbors for UMAP. Since the perplexity is the effective number of neighbors, this makes the runs comparable across both algorithms. For both Fig. S5 (B) and Fig. S6 (B) the rows represent different choices for the parameter (*p* = 2, 5, 15, 30) (*perplexity* for tSNE, *neighbors* for UMAP). For the case of the circle we found that only *perplexity* = 15 we could preserve the *H*_1_ homology class (third row, first column). Furthermore, we found that for all choices of perplexity, there were cases were UMAP could not accurately preserve the topology of the circle. On the other hand, the case of the sphere was more extreme, as both of these algorithms completely vanished the *H*_2_ homology class of the sphere (Fig S6). As a whole there results provide experimental evidence that both UMAP and tSNE could prohibit the discovery of non-trivial topologies in biological datasets.

### 5.4 A set of oscillatory circuits can generate 2− homology

The dynamical system is thus described by the following set of six differential equations:

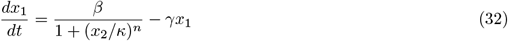

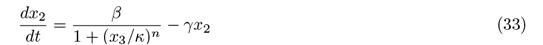

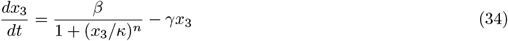

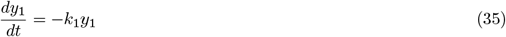

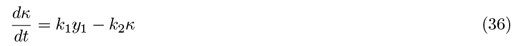

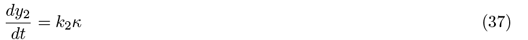

and is thus a modification of the classic repressilator model (51) where the repressor binding strength *κ* is variable. More specifically, the repressor binding strenght dynamics varies as a simple chemical reaction *y*_1_ *→ κ → y*_2_. We performed simulation of this dynamical system and computed its persistent homology. We found that this system contains a persistent 2− homology class and is thus topologically a horn torus (Fig S7).

## 6 Acknowledgements

We thank Alejandro Granados for insightful discussions, Jorge Flores for discussions on algebraic topology and Jesus Rodríguez-Viorato for conversations regarding homology group representatives.

---

**Figure S1:**
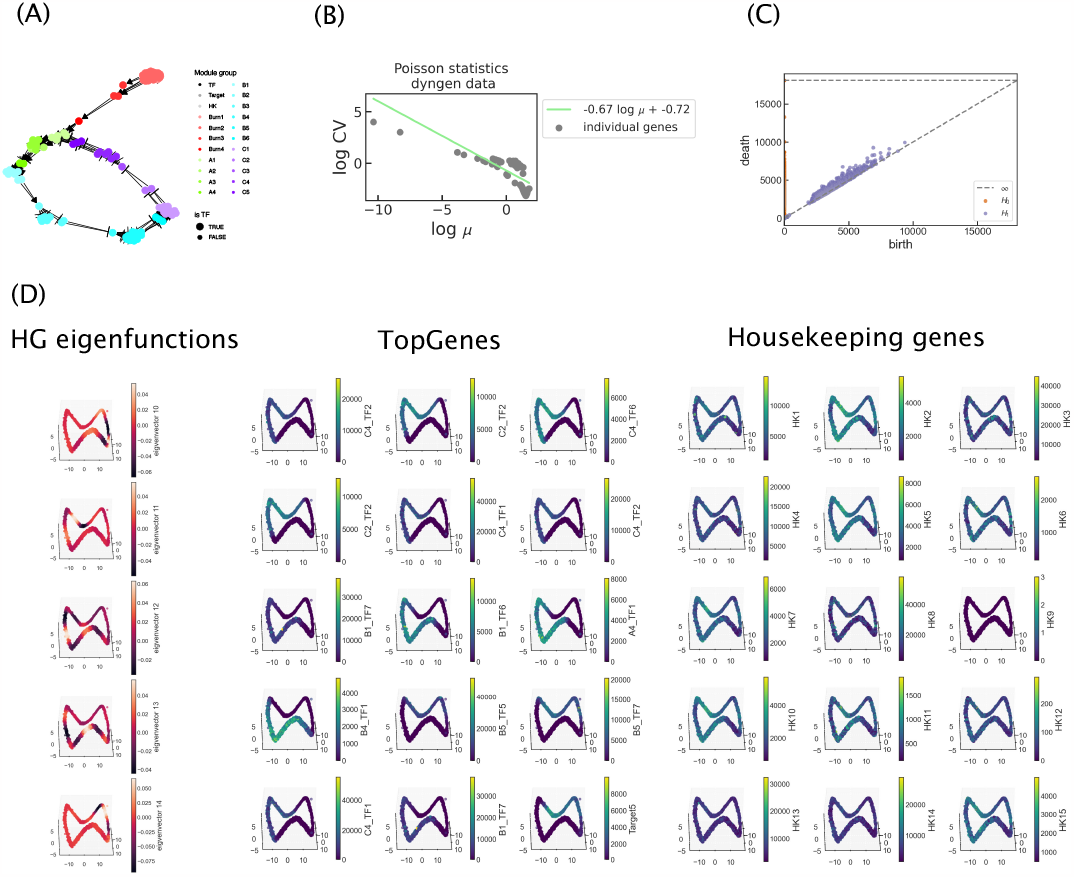
Benchmarking TopGen using a known gene regulatory network. (A) Gene regulatory network used for benchmark experiments. (B) Dyngen data has Poisson statistics. Observed slope of log *μ* vs logCV is − 0.67 which is close to the predicted − 0.5 of Poisson statistics. (C) Persistence diagram using only housekeeping genes. Note that no salient persistent 1− homology classes are present. (D) Left: Eigenvectors of the 0− Laplacian of homology generator. Middle: TopGenes with highest mutual information for the Laplacian eigenvectors on the left. Please note that the gene expression patterns are transient. Right: Examples of housekeeping genes; note that their expression is spurious or constant.

**Figure S2:**
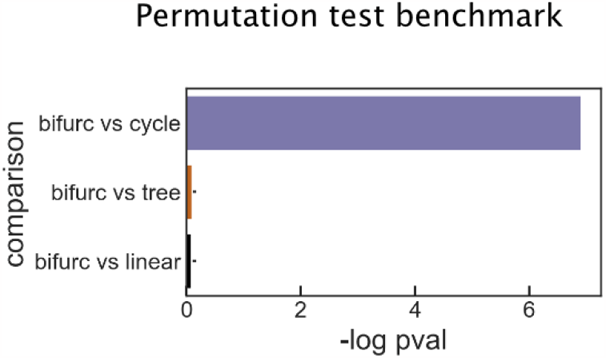
Benchmarks of topological statistic tests using synthetic data. We performed 3 control experiments to evaluate the efficacy of our statistical test. For all experiments we used a simple bifurcation tree as a null hypothesis dataset. For a positive control, we tested a cyclic dataset as a test dataset and found that the difference between maximal lifetime of *H*_1_ classes was significant (P-value *<* 10^− 4^). In contrast a linear, another binary tree, and a trifurcation datasets where all deemed to have a non-significant difference between the maximal *H*_1_ classes (P-values = 0.21, 0.54, 0.4 respectively).

**Figure S3:**
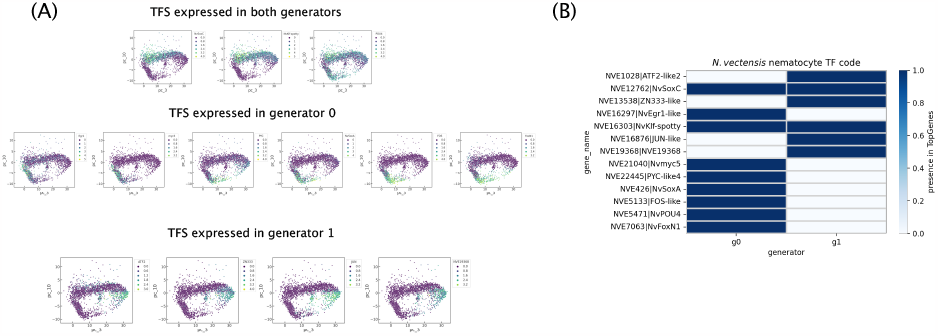
Orthogonal transcription factor code of cnidocytes in *N. vectensis*. (A) Visualization of transcription factor expression along the homology generators. (B) Transcription factors are orthogonally expressed on the homology generators.

**Figure S4:**
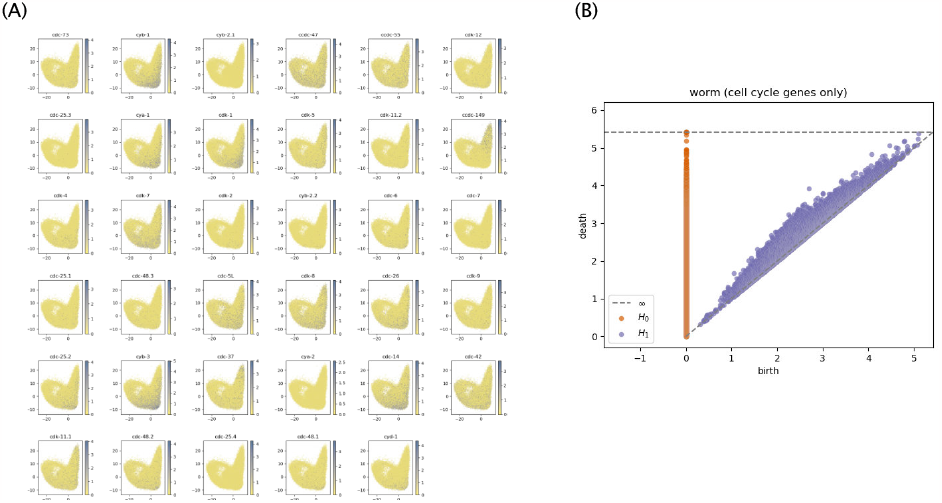
Worm homology generator is not caused by cell cycle. (A) Gene expression of cell cycle genes is spurious along homology generator. (B) Persistent diagram using cell cycle genes displays no persistent *H*_1_ homology class.

**Figure S5:**
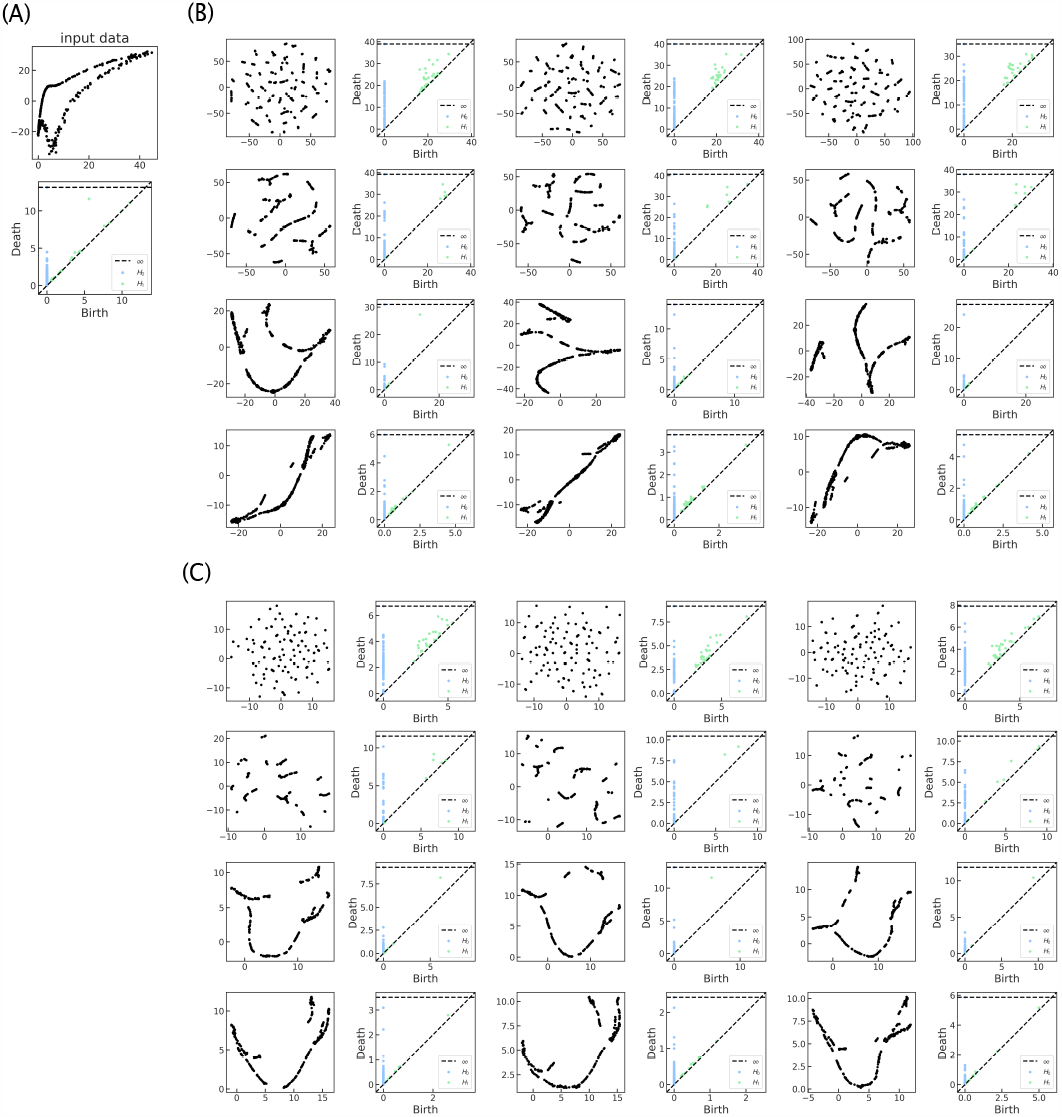
*H*_1_ class can be destroyed using dimensionality reduction methods. (A) The input data consists of a circle after a diffeomorphism (top). Persistence diagram of the morphed circle (bottom). Note the presence of a persistent *H*_1_ homology class (green points). (B) Results using tSNE. The rows represent different choices for the perplexity parameter (*p* = 2, 5, 15, 30). Please note that using *perplexity* = 15 we could preserve the *H*_1_ homology class (third row, first column). (C) Results using UMAP. The rows represent different choices for the number of neighbors parameter (*p* = 2, 5, 15, 30).

**Figure S6:**
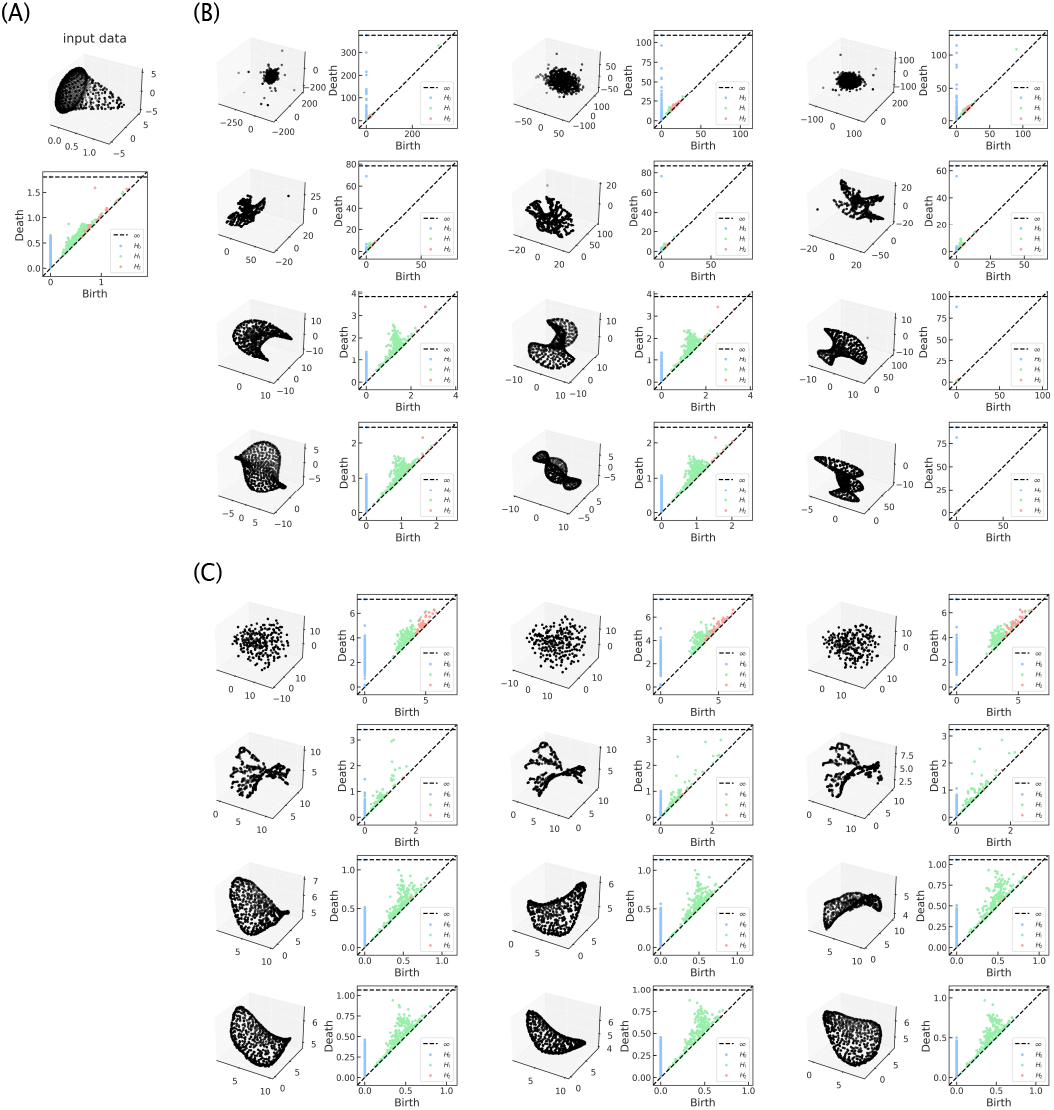
*H*_2_ class can be destroyed using dimensionality reduction methods. (A) The input data consists of a sphere after a diffeomorphism (top). Persistence diagram of the streched sphere (bottom). Note the presence of a persistent *H*_2_ homology class (orange points). (B) Results using tSNE. The rows represent different choices for the *perplexity* parameter (*p* = 2, 5, 15, 30). Note that all replicates fail to preserve the topology of the sphere. (C) Results using UMAP. The rows represent different choices for the *number of neighbors* parameter (*p* = 2, 5, 15, 30). Please note that, similar to tSNE, all replicates fail to preserve the topology of the sphere.

**Figure S7:**
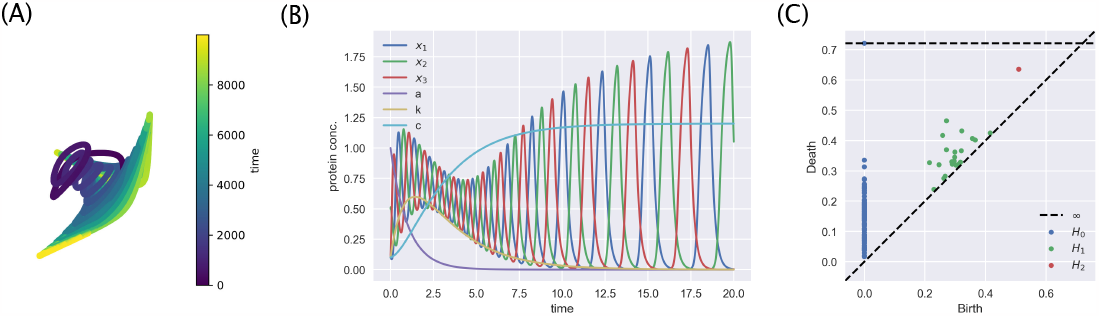
A simple genetic circuit can generate an *H*_2_ class. (A) Phase space of the oscillator proteins in the dynamical system. (B) Time series of dynamical system. (C) Persistence diagram of samples from the phase space. Note that a persistent *H*_2_ class is present (red points).

**Figure S8:**
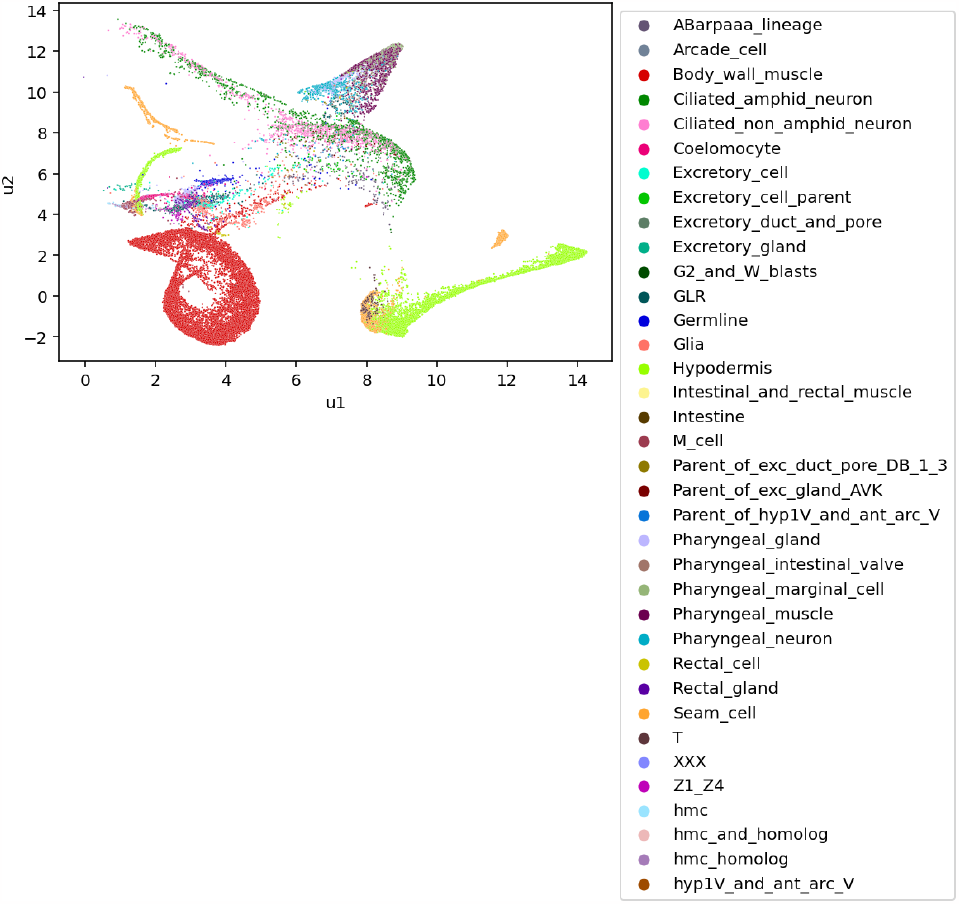
Seam cell loop of *C. elegans* can be destroyed with UMAP.

